# Different Multiword Verb Categories are Processed Differentially in the Brain: An Evidence from EEG Analysis and Decoding

**DOI:** 10.1101/2025.06.06.658278

**Authors:** Hassane Kissane, Nikola Koelbl, Achim Schilling, Patrick Krauss

**Affiliations:** Chair of English Philology and Linguistics, University Erlangen-Nuremberg, Erlangen, Germany; Cognitive Computational Neuroscience Group, University Erlangen-Nuremberg, Erlangen, Germany; Neuroscience Lab, University Hospital Erlangen, Erlangen, Germany; University Hospital Mannheim, Erlangen, Germany

**Keywords:** Multi-Word Verbs, Phrasal Verb, Prepositional Verb, Neural Representation, ERP Analysis, MVPA

## Abstract

The mental representation of multiword verb constructions is a central question in neurolinguistics: are they stored as single lexical units or completely compositional items? This study investigates the neurocognitive processing of two multiword verb types: phrasal verbs (e.g., look up) and prepositional verbs (e.g., decide on). Taking in consideration verb-to-infinitive constructions (e.g., want to go) as a control group. We analyse event-related potentials from eleven native English speakers who completed a listening task while EEG data were recorded. Grand-averaged waveforms and root-mean-square amplitudes were analysed across four-time windows. Therefore, statistical comparisons showed a significantly larger N400 amplitudes for prepositional verbs compared to phrasal verbs, while no significant differences were found between prepositional and to-infinitive constructions. Multivariate pattern analyses confirmed neural discriminability between phrasal and prepositional verbs, but not between prepositional and verb-to-infinitive structures. These results confirm that prepositional and verb-to-infinitive constructions are processed compositionally via valency-based integration, whereas phrasal verbs are stored as lexicalized units. The findings support a theoretical model in which multiword verb constructions differ in their degree of lexicalization, with measurable consequences for real-time neural processing.

## 1 Introduction

Phrasal verbs and prepositional verbs represent two essential multiword constructions in English, blending syntactic complexity with different semantic properties. While both considered verb-particle combinations and multiword constructions (Quirk, Greenbaum, Leech, & Svartvik 1985), the combination is called a phrasal verb when the particle is adverb (e.g., come back, give away), and called a prepositional verb when the particle is a preposition (e.g., agree on, look after). Their processing mechanisms may differ due to their varying degrees of semantic compositionality and syntactic flexibility (Fraser, 1976).

Psycholinguistic and neurolinguistic studies suggest that multiword expressions (MWEs) like phrasal verbs may be stored as single lexical units in the mental lexicon, particularly when they are idiomatic or highly frequent (Arnon & Snider, 2010; Tremblay & Baayen, 2010). However, the neural correlates that underlie the distinction between phrasal and prepositional verbs are still not explored currently.

There was evidence from neuroimaging studies showed that the brain responds differently to stored versus compositional phrases (Van Lancker Sidtis, 2004). Stored linguistic items can mean several types of formulaic expressions such as multiword verbs that may engage similar cognitive mechanisms as other formulaic expressions, which means they might be processed differently than novel constructions. For instance, idiomatic or lexically entrenched expressions often evoke reduced N400 components (Kutas & Hillyard, 1980), related to the word retrieval and the decreased integration process (Kutas & Federmeier, 2011; Lau, Phillips, & Poeppel, 2008; Rommers, Dijkstra & Bastiaansen, 2013). In contrast, compositional constructions -based on their syntactic structure-might raise larger N400 amplitudes, and in cases of syntactic complexity or ambiguity, P600 effects reporting syntactic reanalysis or repair (Kuperberg, 2007; Osterhout & Holcomb, 1992). Furthermore, the semantic P600 reflects difficulty in integrating thematic roles when syntactic and semantic signs conflict (Bornkessel-Schlesewsky & Schlesewsky, 2008).

From a syntactic perspective, phrasal verbs like *give up* or *take off* display non-canonical behaviour. The particles can be separated from the verb (e.g., *take the jacket off*), creating challenges for both syntactic parsing and storage-based models. Prepositional verbs (e.g., *look at* or *rely on)*, in contrast, show stronger syntactic cohesion and often more transparent semantic composition. Despite these differences, both constructions can appear similar on the surface, making them ideal for examining the neurocognitive mechanisms of multiword processing:

1. *She took the jacket off. (phrasal)*
2. *She took off the jacket. (phrasal)*
3. *She looked at the picture. (prepositional)*
4. *\*She looked the picture at. (prepositional)*

A small number of EEG studies have begun to address how the brain processes MWEs, including idioms, binomials, and collocations (Tremblay et al., 2011; Molinaro & Carreiras, 2010; Garibyan et al, 2022) but few have contrasted phrasal and prepositional verbs in a comparison manner. Furthermore, the neural dynamics of how these constructions are syntactically combined and semantically integrated, particularly in real time, are not well understood. Given their lexical and syntactic properties, understanding their neural basis provides strong evidence on the cognitive architecture of language and linguistic theory.

Moreover, recent theories such as Construction Grammar (Langacker, 1987; Fillmore, Kay & O’Connor, 1988; Goldberg, 1995; Croft, 2001; Goldberg, 2005) evidenced by magnetoencephalography (MEG) findings (Cappelle, Shtyrov & Pulvermüller 2010; Pulvermüller, Cappelle & Shtyrov 2013) propose that phrasal verbs are stored as single lexical units with unique form-meaning mappings. This aligns with empirical findings that idiomatic expressions may bypass compositional processing routes under certain conditions (Sprenger, Levelt & Kempen, 2006). Yet, whether prepositional verbs are accessed as single lexical units or compositionally decoded, especially compared to the evidenced unity of phrasal verbs is still debated between different linguistic models (Quirk, Greenbaum & Leech, 1985; Herbst and Schüller, 2008).

### 1.1 Neurophysiological Evidence for the Lexical Unity of Phrasal Verbs

A central issue investigated in the study by Cappelle et al. (2010) is whether phrasal verbs (e.g., *turn up, give in*) are syntactic phrases assembled from independent syntactic elements or lexical units stored holistically in the mental lexicon. The study addressed this debate using MEG to measure early brain responses to particles (e.g., *up, in*) when they appeared in real combinations (e.g., *rise up*) versus pseudo-combinations (e.g., *fall up*). It used a multi-feature mismatch negativity (MMN) paradigm (Näätänen, 1995; Näätänen et al, 2007), which distinguished between lexically stored and syntactically constructed stimuli based on the amplitude of neural responses.

The findings suggested that the particles in real phrasal verbs showed significantly stronger responses than those in pseudo-combinations, beginning as early as ∼190 ms post-stimulus. This lexical entrenchment effect parallels previous MMN findings for morphologically complex words and supports the hypothesis that phrasal verbs are processed as single lexical units, not as syntactic constructions.

More important, this effect was reported regardless of semantic transparency or idiomaticity; both literal combinations (rise up) and metaphorical ones (heat up) showed similarly recorded responses, which challenges any assumptions that only semantically opaque or idiomatic phrasal verbs are lexically stored. Thus, the results align with models of the mental lexicon that allow for redundant storage of frequent compositional sequences and support a lexicalist view of phrasal verbs, wherein phrasal verbs showed differences at early stage of comprehension that aligns with previous studies on early lexical processing (Hauk, Davis, Ford, Pulvermüller & Marslen-Wilson, 2006; Pulvermüller & Shtyrov, 2006; Shtyrov & Pulvermüller, 2007).

### 1.2 Theoretical Background: Valency Theory and the Status of Multiword Verb Constructions

Both the category, phrasal verbs and prepositional verbs, have been analysed in traditional models of grammar as single lexical units (Quirk et al, 1985), which assume that a combination of phrasal verbs (look up) and a combination of prepositional verb (agree with) may be stored in the same way in the mental lexicon. Thus, they should be listed in dictionary as independent Lexemes.

However, a central contribution to this debate comes from the valency theory (Herbst, Heath, Roe & Götz, 2004; Herbst & Uhrig, 2019) a cognitive linguistics and usage-based constructionist model for linguistic analysis, particularly as articulated by Herbst and Schüller (2008), who challenge the traditional treatment of both categories as single lexical units. Instead, they proposed an account grounded in valency-driven complement structures.

The valency theory proposes that verbs possess a set of potential valency slots that specify the number and types of complements a verb may ground. This approach treats phrasal verb combinations not as single lexical entries (e.g., *agree on, write about*), but as regular syntactic structures in which prepositional particles introduce their own complements. For example:

(5) *I wrote to you about it*.
(6) *She worked on the nineteenth-century industrial novel*.
(7) *They were writing about the political solutions*.

In sentence (5), *write* opens two valency slots, each filled by preposition–complement combinations (*to you, about it*). This compositional analysis contrasts with the prepositional-verb approach (Quirk et al. 1985), in which expressions like (e.g., *agree on* or *write about*) are treated as single lexical units requiring separate dictionary entries. Under valency theory, however, *agree* simply requires a preposition (*on*) as a part of the complement not as a verb formation (**Table 1**), which then introduces its own complement:

**Table 1:** Comparison of verb-complement structure analyses in *Quirk et al. (1985)* and *Herbst & Schüller (2008)*. The Quirk et al. model treats combinations such as *agree on* and *agree with* as fixed multiword verb units (prepositional verbs), while the valency-based approach of Herbst & Schüller analyzes the verb as a single lexical item that licenses a complement introduced by a preposition (e.g., *agree* → [on + NP], [with + NP]). This distinction reflects a broader theoretical debate between lexicalist and valency-driven accounts of prepositional verbs.

| Model of Analysis |  |  |  |
| --- | --- | --- | --- |
| Quirk et al. 1985 |  | Herbst and Schüller (2008) |  |
| verb | complement | verb | complement |
| agree | to come early | agree | to come early |
| agree on | a common approach |  | on a common approach |
| agree with | the last bit |  | with the last bit |

(8) *We agreed on the price*.

This syntactic treatment of prepositional verbs differentiates them from phrasal verbs, and alternatively, consider them similar to verb-to-infinitive constructions, such as:

(9) *He agreed to be nominated*.
(10) *She wants to leave early*.

Herbst and Schüller (2008) argue that verb-to-infinitive constructions are structurally parallel to verb-preposition-combinations^1^ constructions: the particle *to* introduces a non-finite verb phrase (*to be nominated, to leave early*) as its complement. Thus, both prepositional verbs and verb-to-infinitive constructions are analysed as valency-driven and compositionally assembled.

By contrast, phrasal verbs such as *look up* or *give in* are often semantically opaque, syntactically discontinuous, and lexicalized, (e.g., *look the number up*). These properties support their treatment as stored single lexical units in the mental lexicon, rather than as transparently decomposed valency structures, which already find experimental evidence as mentioned above (Cappelle et al. (2010).

This study uses event-related potentials (ERPs) to examine the neurocognitive processing of phrasal and prepositional verbs during sentence comprehension and consider verb-to-infinitive as control group for comparison. We compare these verb types under language comprehension task while participants listen to continuous speech. Our goal is to assess whether different ERP components, particularly the N400 (semantic integration) and P600 (syntactic reanalysis or anomaly detection), are modulated by the type of multiword verb construction.

These MEG results from (Cappelle et al, 2010) could provide an important empirical basis for our EEG investigation. If phrasal verbs are stored and accessed as unified lexical items, they should report early and distinct neural response compared to prepositional verbs or to-infinitive constructions, which are hypothesized to be compositional and assembled during processing. Our study builds on this foundation by testing whether similar neurophysiological distinctions arise across these three construction types, thereby expanding the scope of investigation from phrasal to prepositional and infinitival verb structures.

We hypothesize that phrasal verbs, will evoke smaller N400s than prepositional verbs, consistent with storage-based access. On the contrary, prepositional verbs, due to their compositional nature, may show larger N400s and P600s depending on the syntactic configuration and semantic predictability, which might be similar to pure syntactic compositions like verb-to-infinitive sequences. This distinction aims to find how multiword constructions are represented and processed in the brain.

## 2 Materials and Methods

### 2.1 Participants

Participants were 11 (6 females, 5 males) healthy, right-handed, and native speakers of English aged between 26 and 62 years (μ = 42.8, σ = 12.0). Handedness was assessed using the Edinburgh Handedness Inventory (Oldfield, 1971), though detailed laterality index values were not available. All participants reported normal hearing and no history of neurological or psychiatric illness. They gave informed written consent prior to participation and received 50 EUR for the full session as compensation. Ethical approval for the study was granted by the ethics board of the University Hospital Erlangen.

### 2.2 Speech Stimuli and Natural Language Text

As natural language material, we used the novel *Harry Potter and the Philosopher’s Stone* by J. K. Rowling (© Bloomsbury Publishing, 1997), which was released in audiobook format by Pottermore Publishing and narrated by Stephen Fry. The audiobook is publicly available and widely accessible through commercial platforms. For scientific use, we selected approximately 10% of the book, specifically the entire first chapter and part of the second chapter, corresponding to a total of 12 audio segments. These segments ranged from 2:49 to 4:11 minutes in duration (μ = 3:21, σ = 0:24), resulting in a total of approximately 40 minutes of continuous speech. The audio was segmented using *Audacity*, with cuts aligned to paragraph boundaries to preserve discourse-level coherence, though no additional editing was applied to avoid splitting individual sentences or words.

The extracted corpus comprises approximately 5,996-word tokens and presents contemporary, everyday English appropriate for child and adult audiences alike. The book was selected due to its clear narrative style, accessibility in both written and spoken forms, and cultural neutrality. This choice allowed us to present naturalistic, context-rich stimuli while maintaining control over segmentation and listening duration for EEG data acquisition.

### 2.3 Stimulation Protocol

For the auditory presentation, the audio playback was delivered through an 8-channel output sound card at an individually adjusted sound pressure level (SPL) of approximately 30–60 dB, ensuring both intelligibility and participant comfort. The participants fixated on a centrally displayed cross on the screen to minimize ocular artifacts. After each audio block, participants were presented with three multiple-choice comprehension questions related to the preceding audio segment. Responses were collected using a gamepad, with buttons corresponding to predefined answer options (e.g., correct, incorrect, no information). This served to monitor attention and task engagement during listening.

Short breaks of approximately 3 minutes were inserted after Blocks 4 and 8, during which participants could rest. Additionally, brief pauses occurred after each question set, allowing participants to reposition and reduce muscular or postural fatigue. These breaks also served to reduce EEG noise accumulation over time. The entire protocol, including audio presentation, comprehension testing, and breaks, lasted approximately 1.5 hour.

### 2.4 Generation of Trigger Pulses with Forced Alignment

In order to automatically generate trigger pulses for the synchronization of the speech stream with the EEG recordings and to mark the boundaries of words for segmenting the continuous EEG data, forced alignment was applied to the audio and corresponding text data. For this purpose, we used the free web-based service WebMAUS (Schiel, 1999; Kisler et al., 2017), which provides reliable automatic phonetic alignment for German and English speech.

WebMAUS takes as input a waveform file of the audio signal and a plain text file (`.txt` format) containing the corresponding transcription. As output, it provides time-aligned annotations at both the word and phoneme levels, as well as a machine-generated phonetic transcription of the input text. Despite the high reliability of forced alignment algorithms, especially when applied to clearly articulated, professionally recorded speech such as audiobooks, we conducted manual spot checks on randomly selected samples to verify alignment accuracy. There were misalignments observed in these checks in the second audio segment, so we avoided them from the analysis, and alignment precision was consistently within an acceptable margin (typically <10 ms).

For the present study, we used only the word-level boundaries to generate EEG triggers, which were later used to extract time-locked epochs to individual words (the phrasal verb, prepositional verb and verb-to-infinitive). The beginning of each word corresponds to the onset of its first phoneme, and the end time aligns with the final phoneme, ensuring that the two boundary lists can be accurately synchronized if needed.

### 2.5 Stimulus Annotation and Verb Classification

The output of aligned text data with their vocal correspondences were automatically annotated for part-of-speech (POS) using the spaCy library (Honnibal et al, 2020) with the `en_core_web_sm` English language model. The tagging pipeline provided POS tags for each token in the corpus. This automatic annotation was used to identify all verb tokens along with their syntactic complements. Following this automatic tagging, we manually reviewed and verified all verb instances to ensure accuracy and to classify each according to its syntactic construction type. Specifically, each verb was manually categorized into one of three target conditions:

- Phrasal Verbs (e.g., *look up, give in*), where a word tagged as [verb] is followed by another word tagged as [adverb]. In some cases, there was a separation between the verb and the following particle as the case of separation (take [verb] the jacket off [adverb]).
- Prepositional Verbs (e.g., *decide on, agree with*), where a word tagged as [verb] is followed by another word tagged as [preposition].
- Verb-to-Infinitive Constructions (e.g., *want to leave, agree to help*), where the verb followed by a complement introduced by to PoS tagged as [preposition].

Only verbs that were unambiguously classifiable and occurred in clearly disambiguated syntactic contexts such as the verbs where the both conditions are present, as the case of what is called phrasal-prepositional verbs (e.g., Come up with) were excluded from the experimental design.

### 2.6 Pre-processing of EEG Recorded Data

EEG data were pre-processed using the MNE-Python (Gramfort et al, 2013) package (v1.8.0). Raw data were recorded in BrainVision format and loaded with a sampling rate of 1000 Hz. All pre-processing steps were performed in Python 3.10 (Van Rossum, 2009), within a custom environment configured via Google Colab.

#### 2.6.1 Data Loading and Initial Inspection

Raw EEG recordings were imported using `mne.io.read_raw_brainvision` and visually inspected for quality. Channels were filtered and plotted to identify anomalies. A copy of the raw EEG was stored, and a separate copy of the stimulus channel (channel 65) was extracted for event alignment.

#### 2.6.2 Downsampling and Filtering

The EEG signals were downsampled to 250 Hz initially and later to 200 Hz to optimize processing time and reduce computational load. A bandpass filter (1–20 Hz) was applied to remove slow drifts and high-frequency noise, ensuring that ERP components like the N400 and P600 remained intact.

#### 2.6.3 Bad Channel Detection and Interpolation

After that we automatically detected bad channels using a variance-based method. Channels with either zero variance or variance exceeding four times the mean of mid-range variances were flagged. Identified bad channels were interpolated using spherical spline interpolation, and the list of bad channels was reset post-interpolation.

#### 2.6.4 Artifact Correction via Independent Component Analysis ICA

Additionally, to correct for stereotypical artifacts, we applied an (ICA) Independent Component Analysis (Bell & Sejnowski, 1997; Back & Weigend, 1997) using the FastICA method with 30 components. Eye-related components were automatically detected based on correlation with frontal EEG channels (e.g., Fp1). Components exceeding a correlation threshold of 0.2 were marked for exclusion. Additionally, components 0 and 1 were manually excluded. The ICA-cleaned data were reconstructed and saved in `.fif` format.

#### 2.6.5 Stimulus Alignment and Event Annotation

Stimulus alignment was conducted by correlating the downsampled stimulus channel with an external stimulus waveform (audio files). This procedure determined the time lag between the EEG recording and the audio onset. Using this shift, timestamps were created for stimulus onsets (e.g., individual clicks or word onsets from a narrated text). Events were constructed as 3-column NumPy arrays specifying the verb, verb onset according to the audio, and PoS tags (based on previous spaCy tags).

#### 2.6.6 Epoch Extraction and ERP Computation

EEG epochs were extracted around stimulus events from –0.5s to +1s relative to the onset of the target verb stimulus (i.e., phrasal, prepositional, or to-infinitive verb). Baseline correction was applied from –0.5s to 0s, and trials with EEG amplitudes exceeding ±100 µV were rejected. ERPs were computed by averaging across trials per condition (phrasal verb, prepositional verb and verb-to-infinitive). For visualization of ERP waveforms, data were further low-pass filtered at 6 Hz or 8 Hz.

### 2.7 Permutation Test

We performed a non-parametric, permutation-based analysis (Maris & Oostenveld, 2007) to assess differences in EEG responses across the three construction types: phrasal verbs, prepositional verbs, and to-infinitive constructions. The EEG epochs were segmented into four consecutive post-stimulus windows of 250 ms each (0–250 ms, 250–500 ms, 500–750 ms, 750–1000 ms). For each trial and time window, we computed the root-mean-square (RMS) amplitude over all EEG channels: 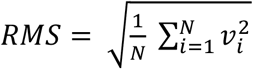 where *v*_*i*_ is the voltage at time point *i*, and N = *f*_*s*_ 250 ms = 50 samples, with a downsampled sampling rate of *f*_*s*_ = 200 *HZ*. This produced a compact summary of signal magnitude per trial and per time window.

To evaluate whether overall RMS amplitude differed between construction types, we first performed a permutation-based one-way ANOVA using 10,000 permutations. RMS data were pooled across participants, and trial labels were randomly permuted to generate a null distribution of F-statistics for each time window. The observed F-values were then compared to this null distribution to derive empirical p-values.

To further investigate condition-specific differences, we conducted pairwise permutation tests between each condition pair (phrasal verbs vs. prepositional verbs, phrasal verbs vs. verb-to-infinitive, prepositional verbs vs. verb-to-infinitive). For each contrast and time window, we computed the observed mean RMS difference and compared it to a null distribution generated by randomly permuting group labels across trials (10,000 permutations). Two-tailed p-values were calculated based on the proportion of permutations yielding a greater or equal absolute difference than the observed one. False discovery rate (FDR) correction was applied across time windows for each pairwise contrast.

### 2.8 EEG-Based Classification using Machine Learning

To investigate whether EEG signals encode condition-specific neural signatures, we conducted three pairwise binary classification analyses comparing each combination of experimental conditions: (1) phrasal verbs vs. prepositional verbs, (2) phrasal verbs vs. verb-to-infinitive, and (3) prepositional verbs vs. verb-to-infinitive.

For each contrast, preprocessed EEG epochs were extracted from all participants and concatenated into a single dataset. Each epoch was labeled according to its construction type. The input feature space consisted of raw EEG signals across all channels and time points, such that each trial was represented as a spatiotemporal matrix (*channels × time samples*).

Before classification, features were standardized using the `Scaler` transformer from MNE-Python and reshaped into 2D format (*samples × features*) via `Vectorizer`. We implemented five commonly used classifiers from the scikit-learn library (Pedregosa et al., 2011): Logistic Regression (Cox, 1958), Support Vector Machine with a linear kernel (Cortes & Vapnik, 1995), Random Forest (Breiman, 2001), Gaussian Naïve Bayes (John & Langley, 1995), and K-Nearest Neighbors (Cover & Hart, 1967). Each classifier was trained and evaluated using 5-fold stratified cross-validation to ensure balanced representation of both classes in training and test splits. Performance metrics—accuracy, precision, recall, and F1-score—were computed at the single-trial level, capturing fine-grained differences in neural response patterns without relying on participant-level averaging.

## 3 Results

### 3.1 ERP Visualization Results

The **Figure 1** displays the grand-averaged ERP waveforms for the three construction types. Overall, all conditions exhibit a similar early sensory response shortly after stimulus onset, followed by a broader centro-parietal negativity that is most pronounced in the 250–500 ms window. In this N400 range, prepositional verbs elicited a significantly more negative deflection compared to phrasal verbs, suggesting increased compositional processing or integration cost. Interestingly, the ERP waveform for prepositional verbs also appears more negative than that of the verb-to-infinitive constructions, although this difference does not reach statistical significance. phrasal verbs and verb-to-infinitive, on the other hand, display more similar ERP profiles throughout the epoch, with overlapping traces in both early and late windows. These patterns are consistent with the hypothesis that prepositional verbs are not stored as single lexical units, unlike phrasal verbs, and are instead processed more compositionally, similar to verb–to-infinitive structures.

**Figure 1:**
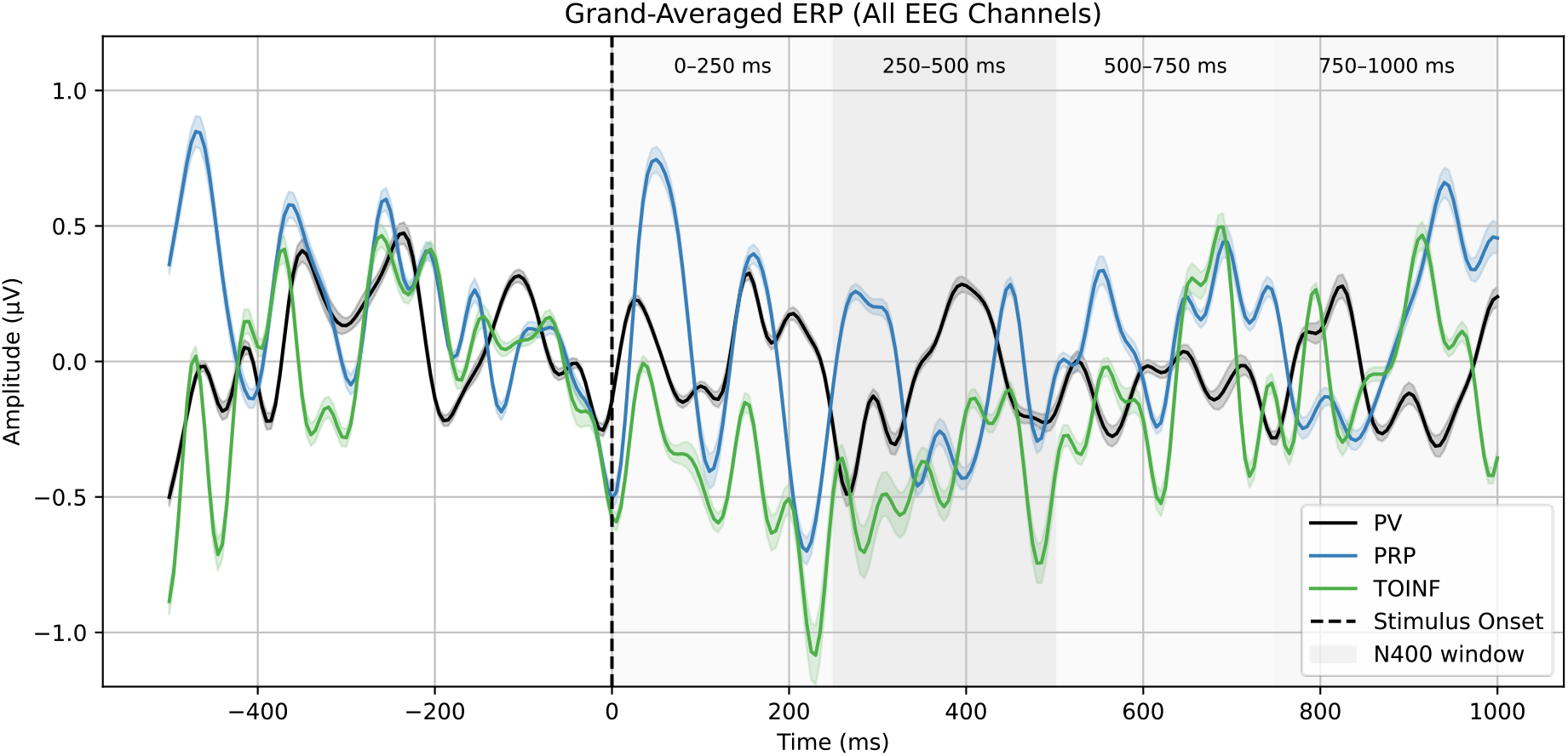
Grand-averaged event-related potentials (ERPs) across all EEG electrodes for the three conditions: phrasal verbs (PV, black), prepositional verbs (PRP, blue), and verb-to-infinitive constructions (TOINF, green). The vertical dashed line at 0 ms marks stimulus onset. Gray background spans indicate the four-time windows used in statistical analysis: 0–250 ms, 250–500 ms, 500–750 ms, and 750–1000 ms. The N400 window (250–500 ms) is additionally highlighted. prepositional verbs evoke a more negative response than phrasal verb in the N400 time window, consistent with reduced lexical integration.

To visualize the spatiotemporal dynamics of ERP responses to the three verb constructions, we generated scalp topographic maps at five key time points: −100 ms (baseline), 0 ms (stimulus onset), 250 ms, 400 ms, and 600 ms. These maps illustrate the evolving distribution of neural activity associated with lexical-semantic and syntactic integration processes.

At 250 ms, a prominent central-parietal negativity emerged in the prepositional verb condition, which intensified around 400 ms. In contrast, the phrasal verb condition exhibited a markedly attenuated negativity at the same time points and scalp locations, indicating reduced semantic integration cost. This contrast aligns with the canonical N400 component, commonly associated with the processing of semantically complex or less predictable structures.

By 600 ms, the phrasal verb condition showed a relative centro-frontal positivity, suggestive of late-stage syntactic reanalysis or integration processes potentially linked to the P600. In comparison, the prepositional verb condition maintained a more posteriorly distributed negativity, lacking the same positivity shift (**Figure 2**).

**Figure 2:**
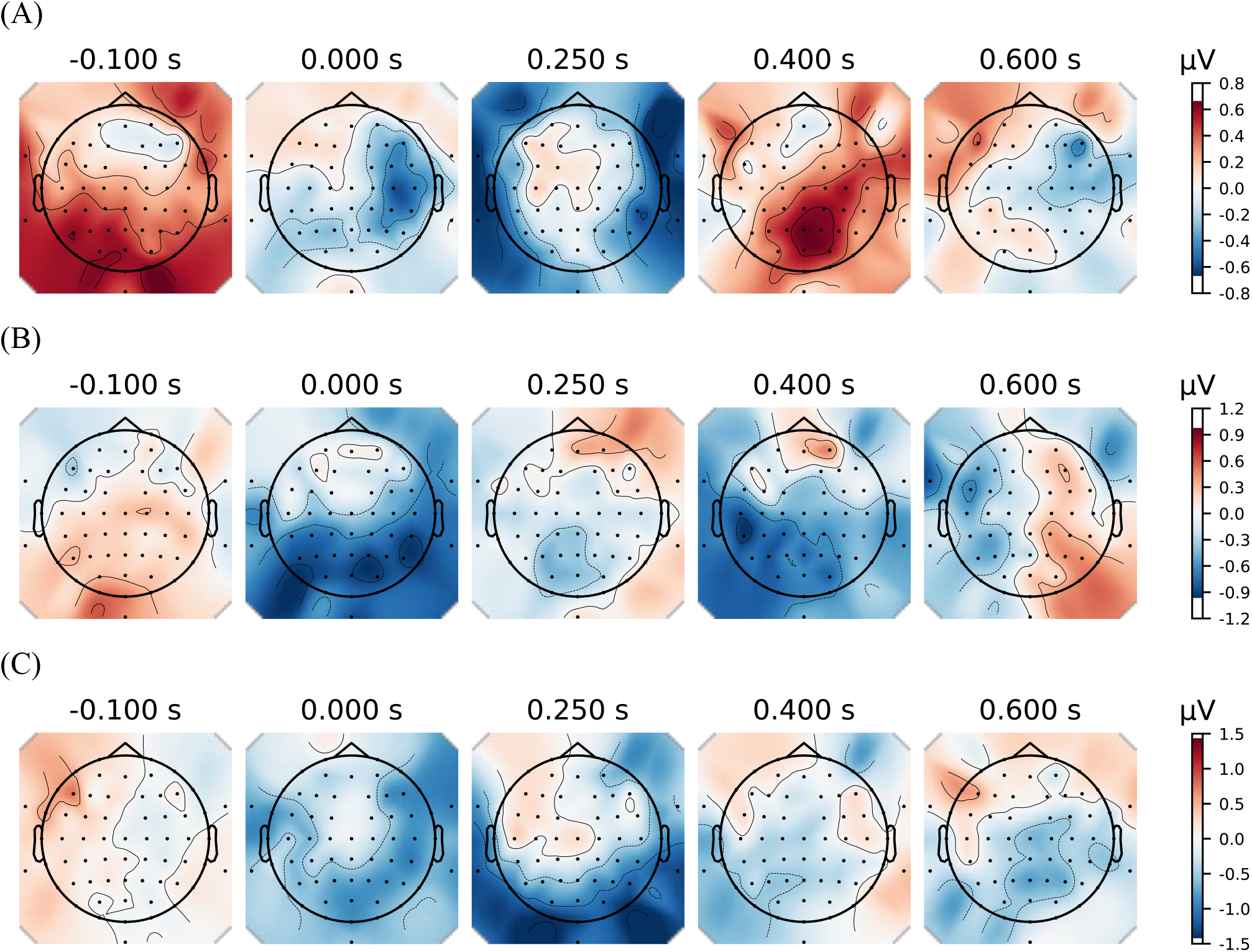
The topographic maps of ERP responses to phrasal verbs (A), prepositional verbs (B), and verb-to-infinitive constructions (C) at five time points relative to stimulus onset (–100 ms to 600 ms). Each map displays the scalp voltage distribution averaged across participants for the respective condition. A pronounced central-parietal negativity is observed in the prepositional verb condition (B) around 400 ms, consistent with the N400 component. Phrasal verbs (A) show reduced negativity in this time window, while the verb-to-infinitive condition (C) shows an intermediate pattern.

To further explore these effects, we computed pairwise difference maps for all condition contrasts (**Figure 3**). The phrasal verb minus prepositional verb maps revealed a robust centro-parietal negativity in the 250–400 ms range, followed by a reversal toward frontal positivity at 600 ms. Although statistical testing did not yield significant clusters after correction, the topographic patterns are consistent with prior ERP research on multiword verb processing and compositional structure. The prepositional verb and verb-to-infinitive conditions displayed more similar patterns throughout, reinforcing the interpretation that they engage overlapping neurocognitive mechanisms.

**Figure 3:**
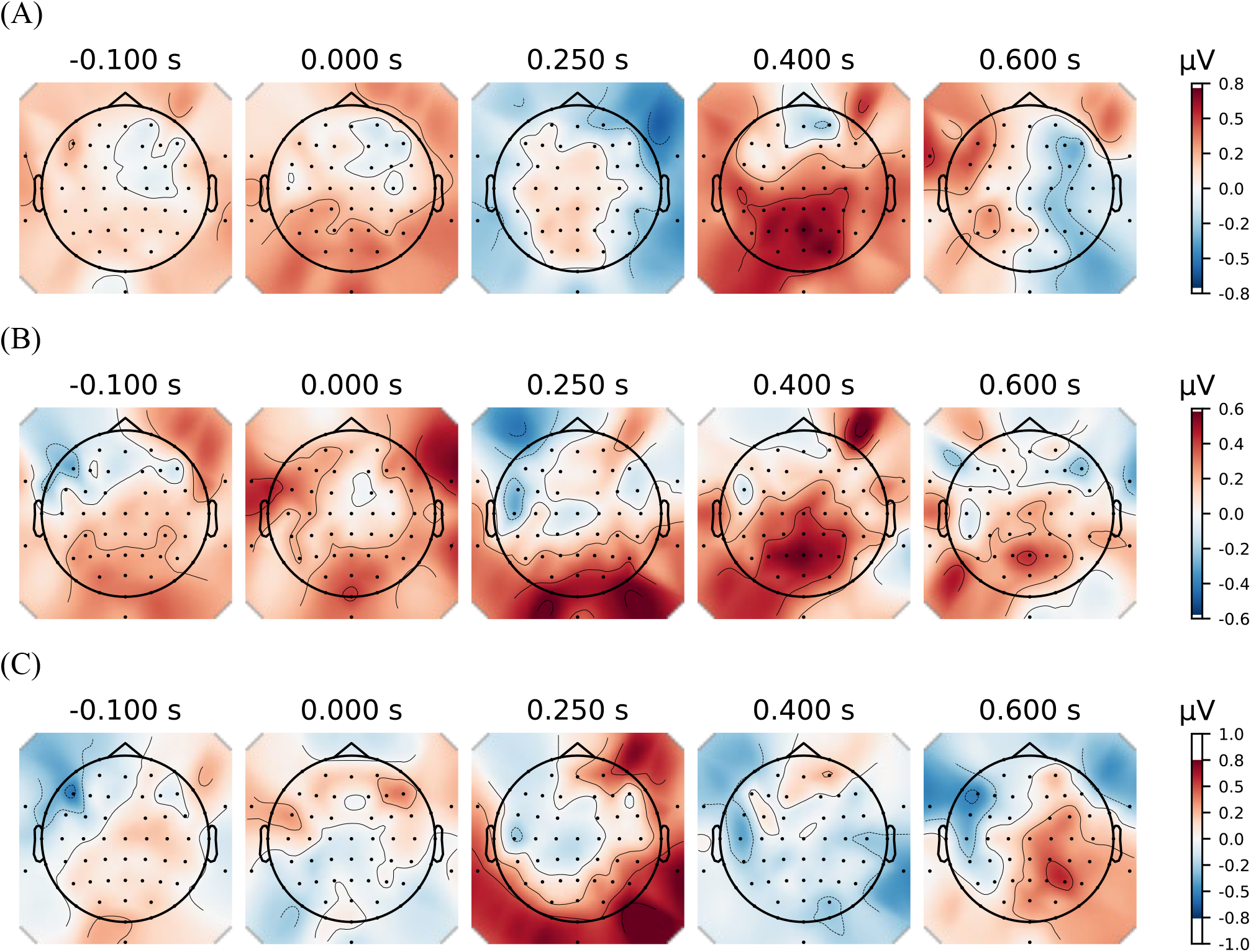
Topographic maps of ERP amplitude differences between conditions at five time points relative to stimulus onset: (A) phrasal verbs vs. prepositional verbs, (B) phrasal verbs vs. verb-to-infinitive constructions, and (C) prepositional verbs vs. verb-to-infinitive constructions. Each map visualizes the scalp distribution of voltage differences, averaged across participants, with red indicating more positive voltage for the first condition and blue indicating more negative voltage. In panel A, a pronounced centro-parietal negativity emerges around 250–400 ms, reflecting a stronger N400 response for prepositional verbs compared to phrasal verbs. The difference between prepositional verbs and verb-to-infinitive (panel C) is more diffuse, with frontal and parietal modulations. The contrast between phrasal verbs and verb-to-infinitive (panel B) shows minimal differences across time points. These patterns support the interpretation that prepositional verbs are processed more compositionally, similarly to verb-to-infinitive constructions, whereas phrasal verbs are processed more like lexicalized units.

### 3.2 RMS-Based Permutation Analysis

To assess condition-specific neural differences over time, we computed the root-mean-square (RMS) amplitudes for each epoch across four consecutive post-stimulus time windows (0–250 ms, 250–500 ms, 500–750 ms, 750–1000 ms) and performed pairwise permutation tests for each contrast. The comparison between phrasal verbs and prepositional verbs revealed significant differences in the two early time windows (**Table 2**): RMS amplitude was significantly lower for prepositional verbs than for phrasal verbs in both the 0–250 ms window (Δ = –0.431 µV, p = 0.0023, p_corr = 0.0086) and the 250–500 ms window (Δ = –0.404 µV, p = 0.0043, p_corr = 0.0086). No significant differences were observed in the later windows (500–1000 ms). In contrast, the comparison between phrasal verbs and verb-to-infinitive constructions showed no significant differences at any time window, with all p_corr values exceeding 0.56. Similarly, the prepositional verbs vs. verb-to-infinitive contrast showed no significant effects after correction, although a numerical trend toward greater negativity for prepositional verbs in the 0–250 ms window was observed (Δ = –0.331 µV, p = 0.0456, p_corr = 0.1824). These findings suggest that prepositional verbs evoke distinct early neural activity compared to phrasal verbs, while their neural profiles are more similar to verb-to-infinitive constructions, consistent with the hypothesis that prepositional verbs and verb-to-infinitive are compositionally processed, in contrast to lexicalized phrasal verbs.

**Table 2:** Results of the RMS-based permutation analysis comparing phrasal verbs, prepositional verbs, and verb-to-infinitive constructions (to-INF) across four post-stimulus time windows. The table reports the difference in mean RMS amplitude (Δ RMS, in µV) between conditions, along with uncorrected (*p*) and FDR-corrected (*p_corr*) p-values. Significant differences after correction are observed in the early windows (0–250 ms, 250–500 ms) for the phrasal verbs vs. prepositional verbs contrast only. No other comparisons reached significance after correction.

| Contrast | Time Window (ms) | $\Delta$ RMS (μV) | $p$ | $p_{corr}$ |
| --- | --- | --- | --- | --- |
| Phrasal vs. Prepositional | 0–250 | –0.431 | 0.0023 | 0.0086 |
|  | 250–500 | –0.404 | 0.0043 | 0.0086 |
|  | 500–750 | –0.142 | 0.2722 | 0.3378 |
|  | 750–1000 | –0.128 | 0.3378 | 0.3378 |
| Phrasal vs. to-INF | 0–250 | +0.100 | 0.5236 | 0.5682 |
|  | 250–500 | +0.208 | 0.1877 | 0.5682 |
|  | 500–750 | +0.096 | 0.5047 | 0.5682 |
|  | 750–1000 | +0.086 | 0.5682 | 0.5682 |
| Prepositional vs. to-INF | 0–250 | –0.331 | 0.0456 | 0.1824 |
|  | 250–500 | –0.197 | 0.2218 | 0.4436 |
|  | 500–750 | –0.046 | 0.7653 | 0.7865 |
|  | 750–1000 | –0.043 | 0.7865 | 0.7865 |

### 3.3 Decoding EEG Group Membership

To evaluate whether EEG patterns differentiate between verb construction types, we conducted pairwise multivariate classification using five standard classifiers: logistic regression, support vector machine (SVM) with a linear kernel, random forest, Gaussian naïve Bayes, and k-nearest neighbors (KNN). Performance was assessed using trial-level accuracy and macro-averaged F1-score to account for potential class imbalance (**Table 3**).

**Table 3:** Classification performance for distinguishing EEG responses across construction types using five machine learning classifiers. Each pairwise comparison (phrasal vs. prepositional, phrasal vs. verb-initive, prepositional vs. verb-to-infinitive) was evaluated using accuracy and macro-averaged F1-score to account for class imbalance. The support vector machine (SVM) showed the best balance of performance in distinguishing phrasal from prepositional verbs, while random forest achieved the highest accuracy across most contrasts but with lower F1-scores. Naïve Bayes and KNN consistently performed near or slightly above chance.

| Classifier | Phrasal vs. Prepositional |  | Phrasal vs. to-Infinitive |  | Prepositional vs. to-Infinitive |  |
| --- | --- | --- | --- | --- | --- | --- |
|  | Accuracy | Macro F1 | Accuracy | Macro F1 | Accuracy | Macro F1 |
| Logistic Regression | 0.61 | 0.60 | 0.58 | 0.56 | 0.55 | 0.55 |
| SVM (Linear) | 0.66 | 0.62 | 0.64 | 0.58 | 0.56 | 0.55 |
| Random Forest | 0.70 | 0.55 | 0.73 | 0.56 | 0.55 | 0.53 |
| Naive Bayes | 0.54 | 0.53 | 0.54 | 0.52 | 0.52 | 0.50 |
| KNN | 0.61 | 0.53 | 0.61 | 0.49 | 0.48 | 0.47 |

For the phrasal verb vs. prepositional verb comparison, the SVM achieved the best overall balance of accuracy (66%) and macro F1-score (0.62). The random forest classifier reached the highest raw accuracy (70%) but had a lower F1-score (0.55), suggesting possible overfitting to one class. Logistic regression and KNN both reached 61% accuracy, while naïve Bayes performed close to chance (54%).

In the phrasal verb vs. verb-to-infinitive contrast, the random forest again yielded the highest accuracy (73%) but with a modest F1-score (0.56). The SVM also performed well (64% accuracy, 0.58 F1), with logistic regression close behind. KNN and naïve Bayes both showed weaker and less stable performance (≤61% accuracy).

For the prepositional vs. verb-to-infinitive classification, overall performance dropped across classifiers. The highest accuracy was obtained by SVM (56%) and logistic regression (55%), with macro F1-scores around 0.55, indicating that these two construction types were the most difficult to distinguish based on EEG patterns.

Together, these results suggest that phrasal verbs are neurally more distinct from both prepositional and verb-to-infinitive constructions, whereas prepositional verb and verb-to-infinitive share more similar neural activation profiles—consistent with the compositional vs. lexicalized hypothesis motivating the study.

## 4 Discussion

This study investigated the neurocognitive processing of two multiword verbs, phrasal verbs, prepositional verbs, in comparison with the control group, verb-to-infinitive constructions, to test whether these structures differ in how compositionally or holistically they are stored and processed in the brain. Based on the theoretical analysis in the valency model (Herbst & Schüller, 2008), we hypothesized that prepositional verbs and verb-to-infinitive constructions would be processed compositionally via valency-driven integration, while phrasal verbs would be retrieved as lexicalized units.

The ERP results support this hypothesis. Prepositional verbs showed significantly greater N400 amplitudes than phrasal verbs in the early post-stimulus windows (0–250 ms, 250–500 ms), suggesting that prepositional verbs require more effortful semantic or structural integration. By contrast, phrasal verbs showed reduced N400 activity, consistent with rapid retrieval from lexical memory. Importantly, no significant ERP differences were observed between prepositional verbs and verb-to-infinitive constructions, reinforcing the idea that both involve compositional interpretation based on valency slots (Herbst & Schüller, 2008). The lack of later (500–1000 ms) differences between any condition pairs further suggests that the critical processing distinctions occur early, likely reflecting differences in lexical access and early structural parsing as for the effects of N400, P600 (Kutas & Federmeier, 2011; Osterhout & Holcomb, 1992) and the early lexical processing at early stages of language comprehension (Hauk, Davis, Ford, Pulvermüller & Marslen-Wilson, 2006; Pulvermüller & Shtyrov, 2006; Shtyrov & Pulvermüller, 2007).

These findings were further confirmed by the machine learning results. The binary classification analyses reported that EEG signals could distinguish phrasal verb from prepositional verb trials, whereas prepositional verbs and verb-to-infinitive trials were more difficult to differentiate. This pattern reinforces the idea that prepositional verbs and verb-to-infinitive constructions engage similar underlying neural representations, while phrasal verbs diverge from both, likely due to their lexicalized nature.

This study contributes both theoretically and methodologically in the study of multiword verbs. Theoretically, our findings align with the valency-based account of multiword expressions and challenge traditional classifications that treat prepositional verbs as lexical units (Quirk et al, 1985). The observed neural similarities between the prepositional verb and verb-to-infinitive suggest that both constructions are processed via syntactic mechanisms rather than through lexical retrieval of fixed expressions. Methodologically, the use of RMS-based permutation testing and multivariate decoding provided converging evidence from both univariate and multivariate neural signatures.

Taken together, our results demonstrate that multiword verbs differ in their degree of lexicalization, with measurable consequences for real-time language processing. Prepositional verbs appear to be more compositionally constructed than traditionally assumed, aligning more closely with verb-to-infinitive structures than with phrasal verbs. These findings support a model of lexical representation that is sensitive to valency structure and compositional depth, rather than relying solely on surface-level form.

Consistent with the valency-based predictions, our ERP results show that prepositional verbs evoke a more negative N400 response than phrasal verbs, suggesting more effortful combinatorial processing. Crucially, no significant ERP differences are observed between prepositional verbs and verb-to-infinitive constructions, supporting the view that both are compositionally processed in real time, whereas phrasal verbs are stored and retrieved as single lexical units.

## Footnotes

1 Therefore, Herbst and Schüller (2008) do not call this combination a *prepositional verb* claiming that this terminology could is used by Quirk et al. 1985 analysis, while they alternatively call them *verb-preposition combinations*.

